# Multi-region spatial transcriptomics reveals region specific differences in response to amyloid beta (Aβ) plaque induced changes in Alzheimer’s Disease (AD)

**DOI:** 10.1101/2025.07.31.667719

**Authors:** Odmaa Bayaraa, Michael Aksu, Evon DeBose-Scarlett, Emily Hocke, Vaibhav Jain, Shih-Hsiu J Wang, Dianne Cruz, Simon Gregory

## Abstract

Alzheimer’s disease (AD) is the leading cause of dementia affecting 55 million people worldwide. The pathological hallmarks of AD, beta-amyloid (Aβ) plaques and neurofibrillary tangles (NFT), follow distinct stereotypical patterns of progression across brain regions and trigger a multicellular response that ultimately leads to neuronal loss and cognitive decline. Despite the uniform spread of Aβ plaque across the cortex during AD progression, different regions demonstrate varying levels of vulnerability and resilience to temporal Aβ plaque induced changes, such as NFT accumulation. There is a critical gap in our understanding of the cell types and molecular mechanisms that underlie these region-specific differences in resilience to Aβ plaque induced changes. In this study, we hypothesized that brain region and cell type specific transcriptional responses within the Aβ microenvironment, and more broadly within the grey matter, may contribute to this variation. We carried out matched multi-region spatial transcriptomics and Aβ immunofluorescence staining from the entorhinal, occipito-temporal, dorsolateral prefrontal and striate cortices from two individuals with Braak III and Thal 4 AD. Spatiotemporal comparisons of cell type proportions, gene expression, and cell-cell communication revealed differences in the vulnerability of somatostatin and somatostatin chondrolectin inhibitory neurons and the expression of endosomal and lysosomal trafficking and metallothionein genes within the Aβ plaque microenvironment. We also observed variations in blood-brain-barrier dysfunction, fibroblast growth factor signaling, and vascular impairment and repair related cell-cell communication networks within the grey matter. Our results demonstrate the value of simultaneously profiling AD-omic and spatial modalities in multiple regions to elucidate how cortical region-specific differences contribute to selective vulnerability and resilience during neurodegeneration.

## Introduction

Alzheimer’s disease (AD), the leading cause of dementia, is characterized by the stereotypical spread of two protein aggregates across the brain: beta-amyloid (Aβ) plaques and phosphorylated Tau (pTau) in the form of neurofibrillary tangles (NFTs)^1-3^. Following the classical amyloid cascade hypothesis, the accumulation of Aβ peptides in the brain has been viewed as the direct cause of AD neurodegeneration^4^, leading to oxidative stress, neuroinflammation, synaptic dysfunction, accelerated NFT build-up, and eventual neuronal loss. However, the limited success of Aβ targeting therapeutics and new insights made possible by advancements in molecular profiling have challenged this framework, pointing to AD neuropathology as a trigger of a more complex “cellular phase” rather than the sole driver of the disease^5,6^. During this cellular phase, which can evolve over decades before clinical symptoms like dementia, the initial cellular interactions and homeostasis initiated by abnormal Aβ and pTau aggregation trigger a chronic downstream response in various brain cell types, leading to AD. This shift has fueled efforts to characterize cellular diversity and function across the different phases of AD progression.

AD disease stage is determined by post-mortem histological staining of Aβ plaque (Thal stages) and NFT (Braak stages) presence across different brain regions. Despite significant evidence of their interplay, the two disease-associated proteins follow distinct patterns of spatiotemporal progression and demonstrate significant differences in brain region vulnerability^7-9^. Technological advancements associated with high resolution single-cell profiling has allowed researchers to refine the individual cell types within brain regions that may be more vulnerable to AD pathology^10-12^. Additionally, developments in spatial transcriptomics and in-situ sequencing have enabled these cell type specific gene expression signals to be interrogated in a native tissue context^13,14^. These studies have demonstrated the importance of studying specific Aβ plaque focal points in a given tissue section, providing an early view into the cellular complexity of the Aβ microenvironment^15-18^.

Although single-cell and spatial studies have provided key insights into plaque-induced gene networks, vulnerable cell subpopulations, and perturbed cellular mechanisms the limitations in cost, tissue availability, and the necessary prioritization of sample size and cohort matching have restricted their application to single brain regions^10,17,19^. Most large-scale human single-nuclei RNA-sequencing (snRNA-seq) and spatial studies have focused on regions of the prefrontal cortex and the middle temporal gyrus, with the exception of Mathys et al 2024^20^. This study used single cell technologies to profile six brain regions from 283 post-mortem brain tissue samples to highlight the role of region-specific gene expression and cell type diversity in AD pathology. Though the snRNA-seq from this study provided critical single cell resolution, the spatial context of these cell type specific responses, especially within and near the Aβ plaque microenvironment, were not described. Therefore, to better understand the contribution of cellular architecture and the molecular mechanisms underlying AD, there is a critical need to profile and compare the Aβ plaque microenvironment from multiple regions within individuals.

The accumulation of amyloid plaques is an early event in AD, and once it triggers the cellular phase, this process leads to downstream pathogenic events including NFT build-up and neuronal death^5,21^. Therefore, we hypothesized that the variable development of NFTs in the entorhinal (ENT-Brodmann area 28), occipito-temporal (OCCP/TEMP-BA37), dorsolateral prefrontal (DLPFC-BA46), and striate (STR-BA17) represent differences in the resilience to Aβ plaque-induced changes in varying regions of the cortex. In this study, we spatially compared gene expression profiles of the Aβ plaque microenvironment between these four cortical brain regions from two AD post-mortem tissue samples. By integrating spatial transcriptomics data with concomitant hematoxylin and eosin (H&E) staining and immunofluorescence imaging of Aβ plaque (6E10) in adjacent sections, we were able to annotate the cortical layers and white matter as well as define ST spots of the Aβ plaque microenvironment within each tissue section. We uncovered differences in cell type composition, gene expression, pathway enrichment, and cell-cell communication between the Aβ plaque microenvironments of the four regions, giving us a unique spatiotemporal view of AD progression and region-specific resilience within the cortex.

## Materials and Methods

### Postmortem human tissue samples

Postmortem brain tissue samples from the two donors were obtained from the Bryan Brain Bank and Biorepository of the Duke and UNC Alzheimer’s Disease Research Center (ADRC)^22^. Informed consent was obtained from all participants and their next-of-kin enrolled in the Autopsy and Brain Donation Program and approved by the Duke Institutional Review Board (IRB number: Pro00110244 and Pro00016278). Cognitive status was determined by annual consensus meetings following contemporary National Institute on Aging and Alzheimer’s Association (NIA-AA) criteria. Post-mortem brain tissue samples were processed and banked according to established protocols^23^. Neuropathological assessment of Alzheimer’s disease neuropathological change was performed according to published guidelines^24^. Subjects with coexisting Lewy body or TDP-43 pathology were excluded.

### Tissue Cryosectioning

The fresh frozen tissue blocks for each cortical region (entorhinal (ENT-BA28), occipito-temporal (OCCP/TEMP-BA37), dorsolateral prefrontal (DLPFC-BA46), and striate (STR-BA17)) were embedded in OCT (Tissue-Tek Sakura) and equilibrated to -20°C in a cryostat chamber prior to sectioning (Leica CM1950). Each block was trimmed until the full surface was exposed and an 8 mm x 8 mm region of interest (ROI) was scored from a region of cortex where all laminar layers of the cortex and white matter were aligned perpendicular to the plane of sectioning. For spatial gene expression permeabilization optimization time, 10um sections from a dorsolateral prefrontal cortex (DLPFC) block ROI were mounted onto the Visium Tissue Optimization Slide, as described in the protocol (10x Genomics, PN-3000394).

From the ROI, 10μm-thick sections were cut and thaw-mounted onto the Visium slides following 10X Genomics Visium guidelines (10x Genomics, #CG000240, Rev E). Briefly, the ROI is placed within the fiducial frame of the Visium Spatial Gene Expression slide (10x Genomics, PN-2000233), thaw-mounted into place by placing a finger on the back of the slide behind the section and thawing the section into place. The slide was then briefly placed on the cryostat peltier to freeze the section and then placed on the cryostat stage to await the placement of the next section. Sections were also taken 10μm serially anterior (two serial sections) and posterior (two serial sections) to the Visium section and thaw-mounted onto charged slides (Thermo Scientific Shandon ColorFrost Plus Slides) and briefly dried on a slide warmer for immunofluorescence staining.

### Visium spatial transcriptomics library preparation

Tissue optimization was carried out following the Visium Spatial Tissue Optimization Reagents Kits User Guide (10x Genomics, #CG000238, Rev C) and 6 minutes was determined to be the optimal permeabilization time. The Visium gene expression slides were then fixed with methanol and stained using Hematoxylin & Eosin (H&E) following the Methanol Fixation, H&E Staining & Imaging for Visium Spatial Protocol (10x Genomics, #CG000160 RevA). The stained slides were imaged using Zeiss Axio Scan Z1 slide scanner using a 20X objective in brightfield. The slides were then used for spatial transcriptomic library construction following the Visium Spatial Gene Expression Reagent Kits User Guide (10x Genomics, #CG000239 RevC), where the tissue was permeabilized for initial cDNA synthesis. Afterwards, second strand synthesis and denaturation were carried out and finally, the cDNA was amplified for library construction. The spatial libraries were then sequenced using an Illumina NovaSeq 6000 platform (Read 1/Spatial Barcode UMI: 28 cycles, i7 index/Sample Index: 10 cycles, i5 index/Sample Index: 10, read 2/Insert: 90 cycles).

### Immunofluorescence staining of adjacent tissue sections

The tissue sections mounted on charged slide were fixed in 1.5% paraformaldehyde (PFA) for 15 minutes and stained with a primary antibody solution (0.25% Triton and 4% BSA) containing mouse anti-β-amyloid antibody 6E10 at 1:10 dilution (BioLegend Catalog # 803014). After overnight incubation at 4°C and three rounds of washes with PBS, a secondary antibody mix containing PBS and donkey anti-mouse IgG conjugated to Alexa 546 secondary antibody diluted at 1:500 (ThermoFisher Catalog # A10036) was added. After a one-hour incubation, five washes with PBS, the slides were cover slipped using the Vectashield (which included DAPI) mounting medium. The slides were imaged using Zeiss Axio Scan Z1 slide scanner at 20X magnification with appropriate filter settings for 360nm and 546nm excitation wavelengths.

### Visium data pre-processing and spot quality control (QC)

10x Genomics Space Ranger software (version 1.1.0) was used to process the raw FASTQ files and H&E .tiff sequencing files. Spot QC metrics for each sample are included in Supplementary Table 1. The Loupe Browser^25^ software’s Visium Manual Alignment wizard tool was used to carry out manual fiducial alignment and identify tissue boundaries. Spots aligning to tissue regions in the H&E that overlap with gaps or folds in the tissue and at the edges of the sections were annotated and removed from downstream analysis. The data from the Filtered feature-barcode matrix, containing only tissue associated barcodes, was used as input into Seurat for further spot and gene filtering.

### Annotation of the Aβ plaque ST spots

Scans from fluorescent imaging were used along with an H&E image of the ST tissue section to annotate Aβ plaque ST spots. Aβ plaque ST spots were identified and annotated when present in adjacent anterior and posterior tissue sections and then visually aligned with the H&E-stained ST section. The labelled H&E image was then used to manually annotate and select ST spots on LoupeBrowser, categorizing the ST spots into three categories: Aβ plaque, Aβ plaque adjacent (2 ST spot boundary around Aβ plaque ST spots) or non-plaque (Justification of the Aβ plaque adjacent is discussed in the results section).

### Quality Control and Filtering of ST spots

Seurat^26^ was used to import and QC the ST raw files. The datasets were loaded in as Seurat objects and their plaque and laminar layer annotations (from Loupe Browser) were added into their metadata fields. Spots that had <10 nFeature_Spatial (total genes per spot) were removed from the dataset. These spots were primarily found in the WM, overlapping with regions predominantly composed of axons, or at the outer boundaries of the tissue section, where cell damage from freezing may be contributing to lower quality ST spots. The percentage of mitochondrial gene expression was calculated for all spots, however, a few spots with high mtDNA % were not filtered out as they also had high nFeature_Spatial counts and may be due to the composition of cells and their metabolic activity^27,28^.

### BayeSpace clustering and laminar layer annotation

BayesSpace clustering was carried out for each sample to guide grey and matter, as well as laminar layer annotations^29^. SpatialCluster() was used with nrep=1000 and q=2 (for grey and white matter annotations) and q=7 (for laminar layers) (bet S1). Referencing the density of cell nuclei, the number of spots determining the cortical thickness of each section, and the boundaries of BayesSpace clusters, grey and white matter, as well as the laminar layers of each section were manually annotated on LoupeBrowser and integrated into the downstream Seurat analysis.

### ST Spot Deconvolution using RCTD

Spot deconvolution was carried out using robust cell type decomposition (RCTD)^30^ from the Spatial-eXpression-R (spacexr) R package^31^. Following quality control and filtering, Seurat objects for each sample were loaded into R along with the Allen brain Human MTG 10x SEA-AD dataset which was converted into a RCTD compatible reference file^32^. The coordinates and count data from each object was used to create SpatialRNA() instances. Next the SpatialRNA instance and reference were used to create an RCTD object and the run.RCTD() function was applied to the objects in ‘full mode’, which does not restrict the number of cells per ST spot. The proportion results were visualized using built-in spacexr functions such as plot_weights() and plot_cond_occur() and ggplot2 package functions. The RCTD weights were also used to plot spatial scatter pie charts using the SPOTlight^33^ package where pie_scale was set to 0.4 to prevent overlap.

### Comparison of Cell Type Proportions

For each sample, the RCTD cell type deconvolution results were then normalized using spacexr’s normalize_weights() function. The WM spots from the Seurat object were then removed to focus our proportion comparisons only on the GM region. Then for each plaque category (Aβ plaque, Aβ plaque-adjacent, non-plaque), the mean weight for each of the 24 cell types was calculated. These mean weights from all samples were then merged into one dataset and visualized using ggplot2^34^ functions (region_plaque_category by cell type matrix). The compiled dataset was converted into a list using the convertDataToList() function with data.type set to ‘proportions’, tansform set to ‘logit’ and scale.fac set to take in the counts of each plaque category for each sample (count of Aβ plaque ST spots in S1_ENT etc.). In model.matrix() we set-up up the design matrix for linear modelling to include region_plaque (ENT_plaque, PFC_non_plaque etc.) as the predictor variable and the sampleID (S1 vs S2) as a covariate. Next, we used the Limma package’s makeContrasts() function to set-up pairwise contrasts for each region (STR_plaque vs STR_non_plaque etc.) and propeller.ttest() was used to compare the proportions of cell types between the groups, with robust set to TRUE and trend set to FALSE. The cell types were grouped by the Common Cell Type Nomenclature (CCN) and were visualized using ggplot2 functions^35^.

### Differential gene expression (DEG) and Gene set enrichment analysis

DEG analysis was carried out using Seurat. For each sample, the raw ST count data filtered to include only GM spots. The data was then used to calculate the percentage expression of each gene and genes that were expressed in at least 1% of spots were retained. Following SCTransform() and clustering, the two samples from each region were integrated using IntegrateLayers() and FindMarkers() was used to carry out pairwise comparisons of gene expression between plaque categories for each region. The results were visualized using functions from the EnhancedVolcano^36^, ggplot2 and ggvenn^37^ packages. DEG results for the contrasts were then used as input into ClusterProfiler^38^ for gene set enrichment analysis. The DEG results were sorted by their avg_log2FC and used as input into gseGO() where all ontologies were included from the human genome annotation database and the minGSSize was set to 50 and pvalueCutoff to 0.05. The Enrichplot^39^ package’s pairwise_termsim() function was used to calculate the similarity between the GO terms and the output was used to visualize the results using the treeplot() function.

### Cell type Specific Cell-Cell Communication Analysis

Comparison of cell-cell signaling networks was carried out using the CellChat package^40^. The two samples from each region were combined to create four region-specific CellChat objects. Each ST spot was assigned a single cell type corresponding to the cell type with the maximum weight for that spot from RCTD spot deconvolution. The receptor-ligand interaction database was created by setting CellChatDB.human and assigning CellChatDB.use a “Secreted Signaling” subset for cell-cell communication analysis. Next, for each region, the communication probability was calculated with type set to “truncatedMean”, trim to 0.1, interaction.range to 250, and contact.range to 100. The results were filtered for min.cells greater than 10 and the cell-cell communication was inferred at a signaling pathway level and aggregated using the computeCommunProbPathway(), where “thresh” was set to the default value of 0.05 to determine a significant interaction, and aggregateNet() functions. Finally, the signaling pathways showing significant communications were further analyzed and visualized for each region using CellChat’s built-in visualization functions such as netVisual_circle() and netVisual_aggregate().

## Results

### Matched multi-region spatial transcriptomics study design

To profile and compare differences in the cellular phase triggered by Aβ plaques, we took advantage of a specific stage of AD progression, choosing two matched samples at Thal phase 4 (Aβ density) and Braak stage III (NFTs). At this stage of AD, Aβ plaques are present throughout the neocortex, but NFTs are concentrated in the transentorhinal region and begin to spread in the temporal and dorsolateral prefrontal cortex but are absent in the striate cortex^8,9^. Hypothesizing that the differential accumulation of NFT pathology reflect a cortical region’s resilience to Aβ induced changes, we decided to profile four representative cortical regions from each sample: entorhinal cortex (ENT, Brodmann Area 28), occipitotemporal cortex (OCCP/TEMP, BA37), dorsolateral prefrontal cortex (DLPFC, BA46) and striate cortex (STR, BA17) (Fig 1A, Table S1).Tissue sections were mounted onto Visium spatial gene expression slides for hematoxylin and eosin (H&E) staining and downstream spatial barcoding with 55um sized uniquely barcoded oligonucleotide spots. Adjacent posterior and anterior tissue sections were used for immunofluorescence staining and imaging of Aβ plaque (6E10).

**Fig 1.**
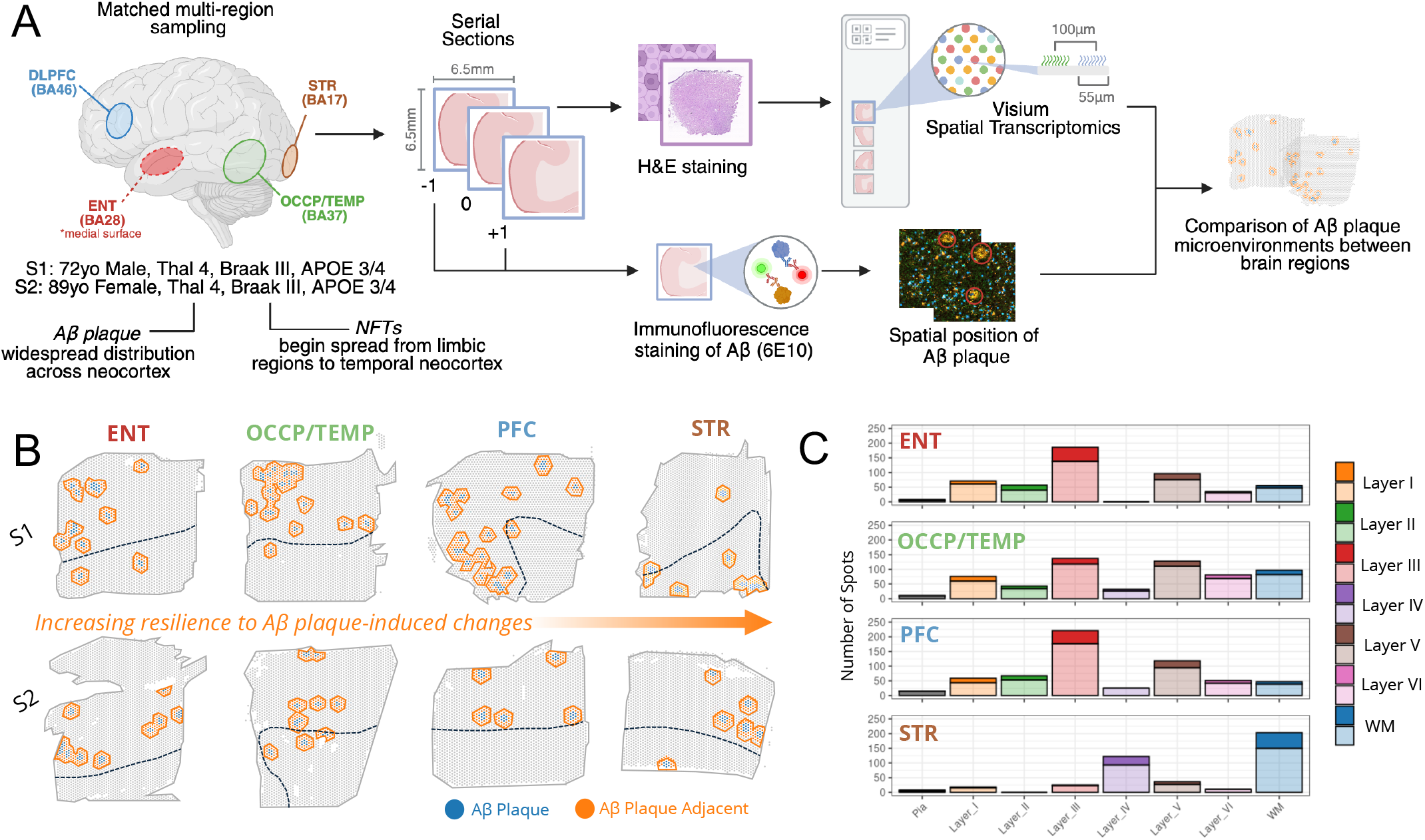
Study Design and Aβ plaque annotation. **A)** Experimental Workflow. Fresh frozen brain tissue samples from two patients were matched for Thal phase, Braak stages and APOE genotype. The ENT, OCCP/TEMP, DLPFC and STR regions of the cortex were selected at Thal phase 4, Braak stage III. From each of the four regions, a 10um thick section was mounted onto a Visium spatial transcriptomics slide and the adjacent serial sections were used for immunofluorescence staining of Aβ plaque. Figure created with BioRender **B)** Annotation of Aβ plaque ST spots. Aβ plaque and Aβ plaque adjacent ST spots were annotated on all tissue sections across the four regions and two samples. **C)** Count of Aβ plaque ST spots across manually annotated laminar layers. The total count of Aβ plaque (darker shades) and Aβ plaque adjacent (lighter shade) spot counts across the manually annotated laminar layers of the cortex.

Adjacent IF stained serial sections were used to annotate the ST spots as either Aβ plaque, Aβ plaque adjacent, or non-plaque (Fig 1B). Aβ plaque adjacent spots were defined as a two ST spot (∼200μm) border around the Aβ plaque annotated spots based on previous evidence that fibrillar plaques range from 30-100μm and diffuse plaques range from 10 to >100μm in size^41,42^. This approach allows us to capture nearby axons, dendrites, and immune cells of the Aβ plaque microenvironment within the constraints of our ST resolution (55 μm spot diameter, 100 μm center-to-center spacing). A combination of H&E imaging and supervised clustering was used to manually annotate the grey (GM) and white matter (WM) boundaries and the laminar layers of each tissue section, where the thickness of specific layers may vary between regions (Fig S1&2). In the ENT, OCCP/TEMP and PFC regions, most Aβ plaque ST spots were found in laminar layers III and V of the cortex, which is consistent with AD histopathology and staging studies^43,44^ (Fig 1C). However, the STR tissue was an exception, as there were a higher number of plaque annotated spots in the layer IV and the white matter^45,46^.

### Regional differences in the enrichment of cell types within the Aβ plaque microenvironment

To determine which cell types are enriched within the Aβ plaque microenvironment, we carried out spot deconvolution of the ST spots using RCTD^30^ (Fig 2A). To maintain consistency of deconvolution across the regions and samples, the Seattle Alzheimer’s Disease Brain Cell Atlas (SEA-AD) single-nuclei RNA-seq dataset, consisting of 166,000 nuclei from fresh frozen middle temporal gyrus (MTG) brain tissue samples, was used to identify proportions of 24 cell types in each ST spot for all samples^47^. To verify the spot deconvolution results, we analyzed the distribution of layer specific cell types in the dataset, such as L2/3 Intratelencephalic Glutamatergic Neurons (L2-3_IT), against our manually annotated laminar layer (Fig 2B). The spots with the highest normalized weights for L2-3_IT cells were mostly found within regions that were manually annotated as either layer II or III, demonstrating concordance between the spatial transcriptomics deconvolution results and manual annotation of laminar layers.

**Fig 2.**
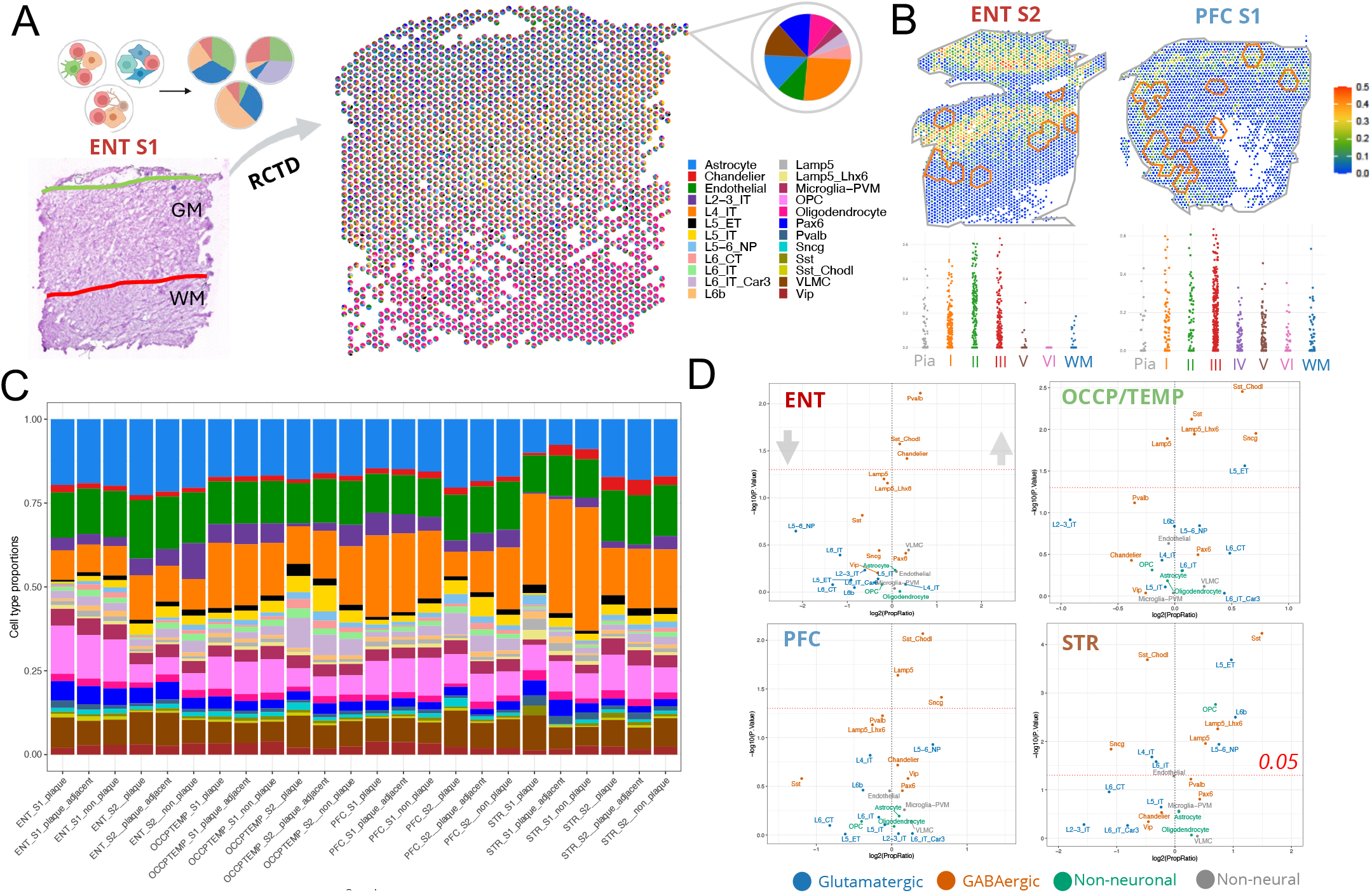
Cell Types Enriched in the Aβ plaque Microenvironment. **A)** Spot Deconvolution of the ST Tissue Sections. RCTD spot deconvolution was carried out using the Allen Brain SEA-AD dataset as a reference to deconvolute the ST spots into proportions of 24 different cell types. Missing spots did not pass the UMI cutoff and are predominantly in the WM of each section, indicative of regions of WM composed of primarily axons, lacking cell bodies. **B)** Proportions of L2/3 Intratelencephalic Glutamatergic Neurons (L2-3_IT) from RCTD spot deconvolution. The normalized weights for each spot (y-axis) plotted against manually annotated laminar layers of the cortex (x axis), indicating that L2-3_IT proportions are highest in manually annotated layers 2,3. **C)** Proportions of Cell Types for Aβ plaque Categories Across all Samples. Barplot colors follow key from Fig 2A. **D)** Differentially Enriched/Depleted Cell Type Proportions the Aβ plaque spots of all regions vs non-pl. Log2FC>0 indicating cell types with higher proportions in Aβ plaque annotated spots, while Log2FC<0 indicate cell types with a lower proportion in Aβ plaque spots. Spots are colored by cell type category.

We then stratified the cell type deconvolution results by three Aβ plaque categories: Aβ plaque, Aβ plaque adjacent, and non-plaque (Fig 2C). To determine which cell types are differentially enriched within Aβ plaque ST spots, we compared the proportions of the 24 cell types between Aβ plaque (not including Aβ plaque adjacent), and non-plaque ST spots for each brain region (Fig 2D). The proportional changes between Aβ plaque adjacent and non-plaque ST spots yielded similar results though with fewer significant cell types as expected (Fig S3). Across the four regions, there was significant enrichment of GABAergic cell types within Aβ plaque ST spots, such as Sst, Lamp5 and Sncg neurons, though with varying magnitudes. Among the GABAergic cell types, the Sst_Chodl interneuron cell type was enriched within the Aβ plaque microenvironment of the ENT, OCCP/TEMP and PFC regions with the exception of the STR (though there is Sst_Chodl enrichment in the STR’s Aβ plaque adjacent spots). Sst_Chodl cells are a rare inhibitory neuron subtype, characterized by their co-expression of somatostatin (Sst) and chondrolectin (Chodl) genes, and have been previously shown to decrease in relative abundance during early AD compared to control in the middle temporal gyrus (MTG)^35,48^. This may be due to differences in the laminar distribution of plaques as Sst_Chodl neurons primarily reside in layer VI and the Aβ plaque spots in the STR were concentrated in layer IV (Fig 1C, S1B). In contrast, there was substantial plaque accumulation in layer VI of the other regions, contributing to the vulnerability of Sst_Chodl neurons. The STR region Aβ plaque ST spots also uniquely demonstrated an enrichment of Sst neurons, several Glutamatergic cell types (L5_ET, L6b, L5-6_NP) and oligodendrocyte progenitor cells (OPCs).

### Comparison of differentially expressed genes (DEGs) and enriched pathways in the Aβ plaque microenvironment between brain regions

We then evaluated the genes and pathways that were differentially expressed and enriched within the Aβ plaque microenvironment. Grey matter spots from the two samples were integrated and pairwise differentially expressed gene (DEGs) analysis performed between different Aβ plaque categories (Fig 3A). The greatest number of significant DEGs (FDR <0.05) were identified for Aβ plaque adjacent vs non-plaque and Aβ plaque vs non-plaque in the OCCP/TEMP and STR regions (Fig 3B). Though most DEGs were unique to each brain region, several genes and gene categories overlapped between the Aβ plaque and Aβ plaque adjacent DEGs (Fig 3C). Notably, glial fibrillary acidic protein (GFAP), a well-established marker gene for astrocytes, and mitochondrially encoded cytochrome C oxidase III (MT-CO3), a mtDNA complex IV gene that demonstrates increased expression in mild cognitive impairment (MCI) and AD blood^49^, were maintained across Aβ plaque and Aβ plaque adjacent vs non-plaque contrasts. The highest number of DEG overlaps (14 DEGs) was observed in the Aβ plaque adjacent vs non-plaque contrast between the OCCP/TEMP and STR regions.

**Fig 3.**
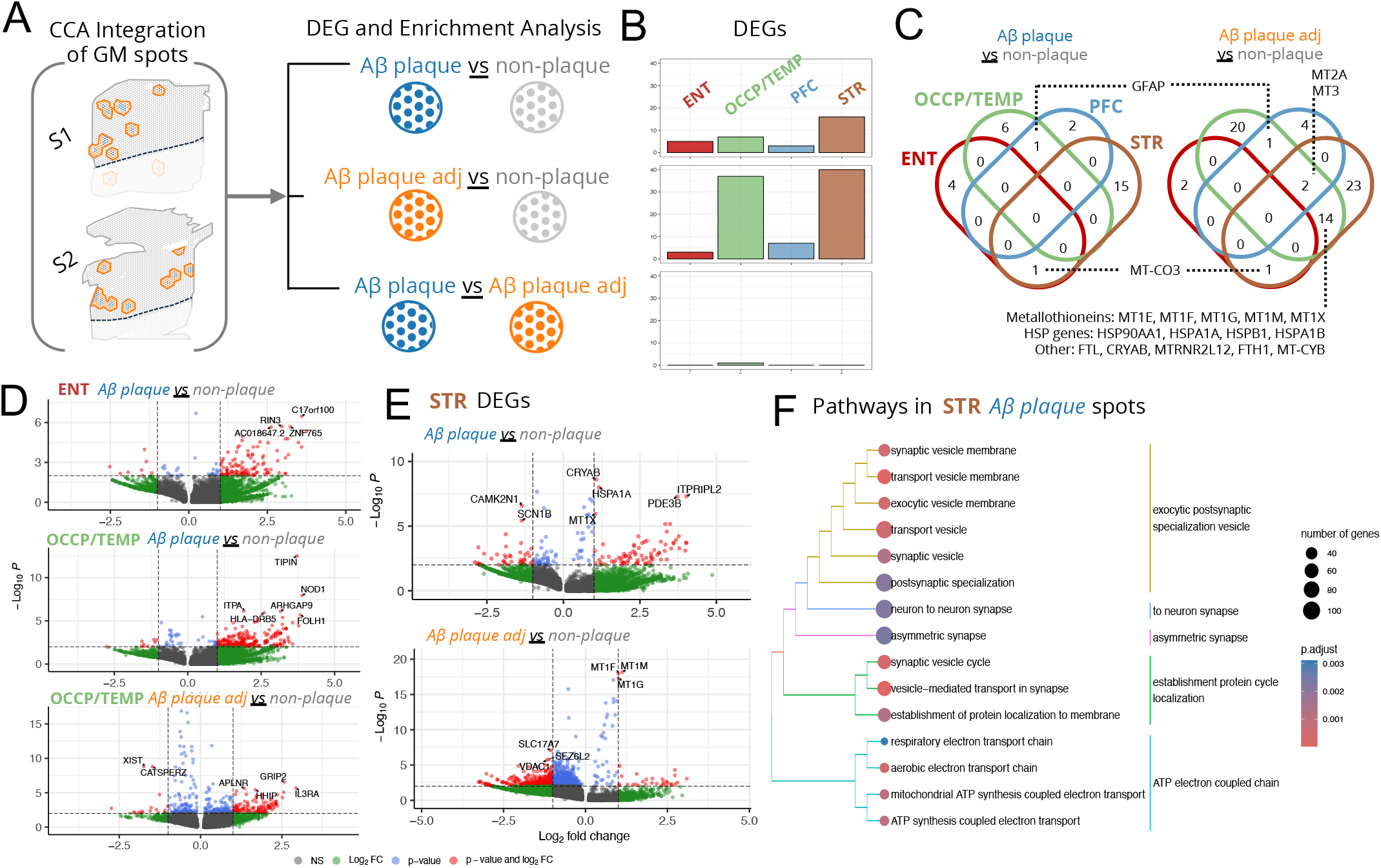
DEGs and Pathway Enrichment in the Aβ plaque Microenvironment. **A)** Differential Gene Expression Analysis Overview. ST data from the grey matter (GM) of each region was integrated across the two samples using CCA integration. Pairwise gene expression comparisons between Aβ plaque, Aβ plaque adjacent and non-plaque ST spots were then carried out. **B)** Number of DEGs for each region for each contrast (FDR < 0.05). **C)** Number of Overlapping DEGs between the 4 regions for each contrast (FDR < 0.05). **D)** Volcano plots of DEGs (FDR <0.05 and abs logFC ≥ 1 are highlighted) in ENT and OCCP/TEMP region Aβ plaque vs non-plaque and Aβ plaque adjacent vs non-plaque contrasts. **E)** Volcano plots of DEGs (FDR <0.05 and abs logFC ≥ 1 are highlighted) in STR. All volcano plots are plotted using nominal p-values for visualization purposes, but only highlighted significant genes with FDR <0.05 and abs logFC ≥ 1 are further discussed **F)** Treeplot of Top 15 Enriched Pathways (p-value <0.05) in the Aβ plaque Microenvironment of the STR region.

AD and neurodegeneration-associated DEGs specific to the Aβ plaque microenvironment of each region were identified (Fig 3D). In the ENT region, the top DEGs in Aβ plaque ST spots included the upregulation of Ras and Rab interactor 3 (RIN3), a late-onset AD (LOAD) GWAS loci that translates into a guanine nucleotide exchange protein involved in pTau and Aβ regulation and a potential biomarker for the disease^50,51^. In the OCCP/TEMP region, ST Aβ plaque spots showed differential upregulation of Nucleotide-binding oligomerization domain (NOD1) gene (also known as NLRC1), an innate immunity gene that yields receptors involved in the regulation of microglia-driven neuroinflammation^52,53^. Additionally, the Interleukin 3 receptor subunit alpha (IL3RA), a key receptor for IL-3 which mediates astrocyte-microglial cross talk in AD^54,55^, was upregulated in OCCP/TEMP Aβ plaque adjacent spots. The STR region contained the most significant DEGs across the Aβ plaque and Aβ plaque adjacent comparisons (Fig 3E). We identified upregulation of alpha-crystallin B chain (CRYAB), a gene implicated in AD which has been shown to prevent astrocyte apoptosis and prevent inflammation^56,57^, and a cluster of metallothionein genes (MT1M, MT1F and MT1G), a class of proteins involved in heavy metal homeostasis and regulation of oxidative stress that have been shown to be upregulated near Aβ plaque in AD mouse and rat cortical models^58,59^.

Gene set enrichment analysis (GSEA) was then carried out using the log fold change values (logFC) from the DEG analysis results. In the ENT region, DEGs between the Aβ plaque and non-plaque ST spots were enriched for several immune related pathways: myeloid leukocyte activation, immune response-activating cell surface receptor signaling pathway, adaptive immune response and immune effector process (Fig S4). The involvement of immune signaling pathways, specifically of myeloid cells, in the ENT is consistent with other AD studies^60,61^. In the STR region, DEGs between the Aβ plaque and non-plaque ST spots identified significant enrichment of pathways involved in exocytic postsynaptic specialization vesicle, neuron to neuron and asymmetric synapse, establishment of protein cycle localization, and the ATP electron coupled chain (Fig 3F). Finally, we identified elevated expression of Extracellular vesicle pathways, which have previously been associated with AD pathogenesis. However, the mechanisms and function are unclear as they have been shown to both carry and spread pathogenic proteins while providing protective effects through growth factors, neurogenesis and clearance of the cellular environment^62,63^.

### Region and cell type specific ligand-receptor interactions

We then analyzed how cell type specific receptor-ligand interactions compared between the four brain regions. We detected 14 signaling pathways where at least one pair of cell types had a significant ligand-receptor interaction (p-value <0.05) across the four regions (Fig 4A). Of these pathways, five pathways were detected in all four regions, including Cyclophilin A (CypA), Prosaposin (PSAP), Pleiotrophin (PTN), Fibroblast growth factor (FGF) and Growth arrest-specific (GAS). Three pathways were detected in three regions, involving Vascular endothelial growth factor (VEGF), granulin (GRN) and SLIT- and NTRK-like (SLITRK). Broadly, these 14 pathways can be grouped into four categories based on their biological function: neurotrophic signaling and growth factors, lysosomal function, inflammation and immune response, and synaptic maintenance and circuit modulation. The inferred intercellular communications networks of these pathways allowed us to compare cell type specific autocrine and paracrine signaling relationships (Fig 4B & 4C, S5). Across pathways detected in all four or three of the four regions, we observed that astrocytes play prominent sender (source of signal), receiver (of incoming signals), mediator (ability to gatekeep the communication of the signaling pathway-flow betweenness) and influencer roles (ability to influence information flow-information centrality) (Fig S5)^40^. In the less resilient brain regions like the ENT, the importance of this role in astrocytes is more prominent and as the resilience to Aβ plaque-induced changes increases into the STR, this signal dissipates.

**Fig 4.**
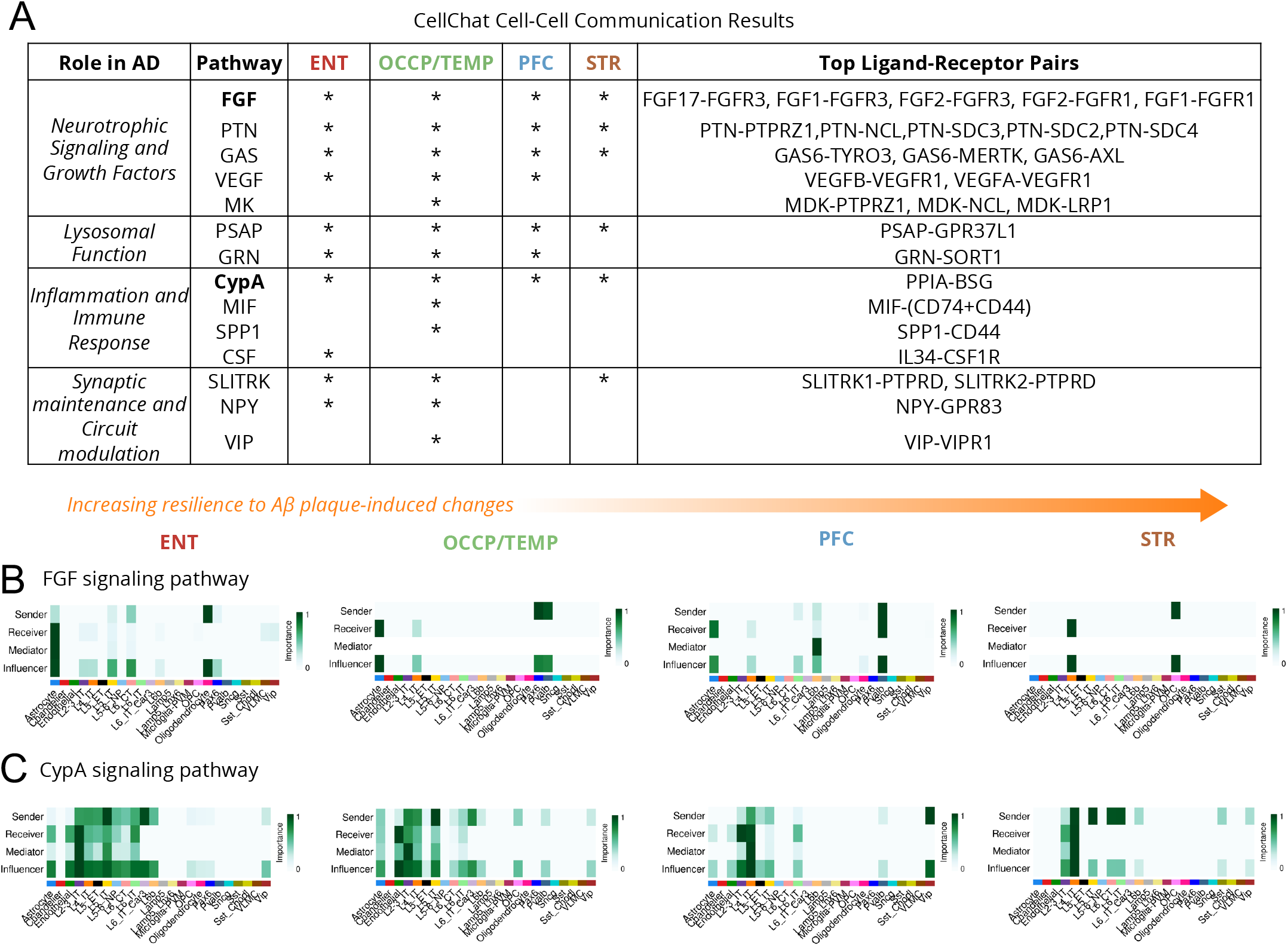
Cell-cell communication networks for different pathways across the ENT, OCCP/TEMP, PFC and STR regions. **A)** Table of All Detected CellChat Pathways. * indicates that at least one significant receptor-ligand interaction was detected between two cell types (p-value <0.05). Heatmap of the expected role for each cell type calculated from the network analysis scores for **B)** FGF signaling and **C)** CypA signaling pathways.

## Discussion

The advent of spatial and single-cell transcriptomic approaches have been crucial for deciphering the cell type molecular mechanisms underpinning AD. Despite useful insights from previous studies using these approaches, their primary application to single brain regions provide a limited view of the differences in selective vulnerability and resilience to Aβ plaque induced changes between regions of the neocortex. Here, we generated spatial transcriptomic data of two AD patients from four brain cortical regions, including ENT, OCCP/TEMP, PFC and STR, representing the different levels of resilience to Aβ plaque induced changes. This matched analysis of cell type proportions, gene expression within the Aβ plaque microenvironments, and cell-cell communication enabled us to compare the cellular and transcriptomic landscapes at the focal points of Aβ plaque accumulation, offering insights into how differences in the cytoarchitectures of these regions may drive varying vulnerability and resilience.

Spatial transcriptomic analysis of cell type composition within Aβ plaque microenvironments revealed that GABAergic cell types, specifically Sst_Chodl neurons, were found in significantly higher proportions in Aβ plaques. The co-localization of Sst- and Sst-inhibtory neurons within Aβ plaque aggregates, as well as their progressive and specific loss in AD, is well characterized^48,64^. SST expressing neurons have been shown to regulate Aβ plaque catabolism through both direct and indirect pathways, contributing to disease pathogenesis^65,66^. Consens *et al*. has demonstrated that there is a robust decrease in Sst interneurons across six different regions of the neocortex, though with varying effect sizes and significance levels^67^. Gabitto *et al*. also found that Sst interneurons decreased with disease progression and demonstrated that vulnerable Sst subpopulations underwent specific molecular changes during early AD, such as the downregulation of genes including nerve growth factor and metalloendopeptidase^47^. Our results, while supporting existing findings on the selective vulnerability of Sst cell types, further highlight Sst_Chodl subtype vulnerability and how these cells may be focally impacted within the Aβ plaque microenvironment. Furthermore, Sst_Chodl neurons are functionally unique long range inhibitory subtype that have been shown to regulate the synchronization of neuronal activity across different brain regions^68^. Additional studies are therefore needed to characterize the proportions of Sst and Sst_Chodl across the cortex and understand how their heterogeneity may quantitatively, through cell death, and qualitatively, through cell-type specific circuits or molecular pathways, contribute to regional selective vulnerability and resiliance^69^.

When we compared gene expression in Aβ plaque ST spots with non-plaque ST spots for each region, there was a notable upregulation of Rab Interactor 3 (RIN3) in the ENT plaque ST spots, a well-established Alzheimer’s disease risk allele^70^. RIN3 has been shown to correlate with endosomal dysfunction in APP/PS1 transgenic mice^71^, a widely used transgenic mouse model of AD. RIN3 also interacts with BIN1, a canonical AD risk gene, to regulate Aβ production and current models hypothesize that its upregulation leads to Rab5 hyperactivation, causing impaired trophic signaling and neurodegeneration^72-74^. From our cell-cell communication results, PSAP and GRN pathways involved in lysosomal function were detected across all four regions (Fig S5). In addition to cell type specific differences in autocrine and paracrine signaling, the GRN pathway was detected in three of the four regions except for the more NFT resilient STR. PSAP and GRN protein levels have been shown to be positively correlated with Aβ and specifically GRN levels with pTau which is consistent with our results^75,76^. This supports studies that suggest differences in endosomal trafficking and lysosomal function and a region’s ability to repair or compensate for its disruptions that induce synaptic loss, which may be an important factor contributing to differences in regional resilience during AD pathogenesis^77,78^. In our dataset, the more NFT resilient STR region demonstrated a unique upregulation of metallothionein genes, where MT1F, MT1M and MT1G were significantly upregulated in the plaque ST spots. On the other hand, MT2A and MT3 were downregulated in the less resilient OCCP/TEMP’s Aβ plaque adjacent ST spots. MT-1/2 upregulation has been consistently observed in AD^58^, and studies comparing AD and resilient individuals suggest that increased MT-1/2 expression in resilient individuals decreases the amount of reactive oxygen species and downstream Aβ oligomers and pTau^79^. Our data further suggests that even within the same individual, there may be variability in metallothionein expression within the Aβ microenvironment of different cortical regions. This may provide more resilience through metal cellular homeostasis and a protective effect against neurotoxic Aβ oligomers, which may confer region specific resilience to pathology.

From our cell-cell communication analysis, Fibroblast growth factor (FGF), a well-established pathway implicated in AD, was detected in all four regions (Fig 4B)^80^. In the ENT, OCCP/TEMP and PFC regions, astrocytes play important receiver and influencer roles, but in the STR, L4_IT and OPC cell types take on these roles, which may be driven by the more prominent L4 in STR cortex. FGFR3 and its ligand FGF2, a common top receptor-ligand pair detected in our dataset, have been shown to regulate extracellular tau internalization in the presence of Aβ^81^. FGF1 and FGF2 have been shown to reduce inflammation, promote neurogenesis and inhibit neurotoxicity among many other neuroprotective functions during AD pathogenesis^82^. These differences in FGF interactions between regions and cell types detected in our dataset could provide novel drug targets considering FGF ligands and receptors are currently being studied extensively for their therapeutic potential in AD^83^. Our pathway analysis also detected Cyclophilin A (CypA), a proinflammatory cytokine that drives vascular impairment, in all four regions (Fig 4C)^84^. CypA has been shown to directly bind with tau and studies have shown that it is elevated in ApoE4 astrocytes leading to CypA-NFkB-MMP-9-dependent blood-brain barrier (BBB) dysfunction^85^. In our dataset, astrocytes play significant receiving and influencing role in the CypA signaling network in the ENT and OCCP/TEMP regions. However, this is not the case in the more resilient PFC and STR regions. Additionally, vascular endothelial growth factor (VEGF) signaling, a key pathway responsible for angiogenesis and vascular maintenance, was identified as significant communication in the ENT, OCCP/TEMP and PFC regions. Unlike their role in CypA, astrocytes primarily send the signal for the VEGF pathway in these regions (Fig S5). Though VEGF is strongly associated with AD, it exhibits complex variable roles and potential impairments during AD pathogenesis that remain to be clearly elucidated^86,87^. Taken together, our results suggest regional differences in BBB integrity and vascular impairment during AD that may be coordinated by the complex dual role of astrocytes that contributes to CypA mediated damage but positively directs VEGF repair and compensation of vasculature. Large scale single-nuclei studies have also demonstrated regional differences in astrocyte transcriptional identity, vascular specialization, the loss of region-specific vascular features, and modifications of neurons and glial cell communication during AD^88,89^.

While our study provides a critical insight into region- and cytoarchitecture specific changes in gene and pathway expression in the context of AB plaque-related pathology there were some limitations. This includes our small samples size and lower resolution ST approach. Additionally, we did not include IF staining of phosphorylated tau (pTau) and we lack the pairing of age, sex, and APOE genotype matched cases and cognitively normal controls as would be carried out by a more expansive study. This design would allow for a better understanding of the complex interactions between the two pathological hallmarks and the region-specific cellular responses to pTau. With new higher resolution ST approaches, made possible by smaller feature and microarray-based capture approaches, the deconvolution of cell types would be mitigated to achieve a true single cell resolution^90^. Further, the inclusion of immunofluorescence staining of proteins on the same tissue section would enable more accurate annotation of pathological features^18,91^.

## Conclusions

In this study, we have demonstrated the value of multi-region ST approaches to better understand the spatiotemporal cellular mechanisms underlying region-specific differences in resilience to Aβ plaque induced changes during AD. Between the ENT, OCCP/TEMP, PFC and STR cortical regions, we compared cell type proportions and gene expression within the Aβ plaque microenvironment and cell-cell communication networks in the grey matter. We found differences in Sst inhibitor neuron vulnerability within Aβ plaque microenvironment, particularly within the Sst_Chodl subpopulation. In addition, we identified differential expression of endosomal and lysosomal trafficking and metallothionein genes that may contribute to mechanisms of resilience to plaque induced changes. Finally, we observed differences in in BBB dysfunction, FGF signaling and vascular impairment and repair within the grey matter that may underlie the vulnerability of specific cortical regions to AD pathology. Our work highlights a key gap by including the analysis of striate cortex as a pathoetiological resilient region for AD research. Future multi-region comparative studies expanded to include additional multi-omics will help us to better understand the regional differences in vulnerability and resilience, which will provide key clues to the underlying mechanisms driving AD.

## Declarations

### Ethics approval and consent to participate

Postmortem brain tissue samples were obtained by the Bryan Brain Bank and Biorepository of the Duke-UNC Alzheimer’s Disease Research Center (ADRC). The Autopsy and Brain Donation Program was approved by the Duke Institutional Review Board (IRB number: Pro00110244 and Pro00016278).

### Availability of data and materials

The raw sequencing data and the Space Ranger processed output files have been deposited in the NCBI Gene Expression Omnibus (GEO) and will be publicly available as of the date of publication.

### Competing interests

The authors declare that they have no competing interests.

### Funding

This work was funded by the Duke and University of North Carolina Chapel Hill (UNC) Alzheimer’s Disease Research Center (ADRC) supported by the National Institute on Aging (NIA) of the National Institutes of Health (NIH) under award P30AG072958. This work was also made possible by support from the Margaret Harris and David Silverman Endowed Professorship at Duke University.

### Authors’ Contributions

Project conceptualization S.-H.W., D.C and S.G.G., Sample preparation and data curation, S.-H.W., D.C, and E.H, Computational analysis and data visualization, O.B, M.A, E.S and V.J, Interpretation and original draft, O.B, S.-H.W., D.C and S.G.G., All authors reviewed and edited the final manuscript.

## Acknowledgements

We would like to thank the Molecular Genomics Core at the Duke Molecular Physiology Institute, Duke University School of Medicine, for their support generating and analyzing the data for this manuscript, especially Karen Abramson and Stephanie Arvai. We also thank the Duke Bryan Brain Bank and Biorepository and Neuropathology core, especially John Ervin. Most importantly, we thank the patients and their families who donated their tissue to the Bryan Brain Bank without whom this study would not have been possible.

## Supplementary Figures and Tables

Supplementary Figure 1

• BayesSpace clustering and annotation of cortex laminar layers for each sample

Supplementary Figure 2

• Spatial expression of layer specific marker genes

Supplementary Figure 3

• Cell types enriched in Aβ plaque-adjacent spots vs non-plaque

Supplementary Figure 4

• Dotplot of GSEA pathways enriched in ENT plaque vs non-plaque DEGs

Supplementary Figure 5

• Additional cell-cell communications networks from CellChat analysis

Supplementary Table 1

• Sample information and Visium spatial transcriptomics raw QC metrics

Supplementary Table 2

• Mean RCTD weights of 24 cell types for each plaque category (Aβ plaque, Aβ plaque-adjacent, non-plaque)

Supplementary Table 3

• List of differentially expressed genes (DEGs) from pairwise comparisons of Aβ plaque, Aβ plaque-adjacent, non-plaque plaque categories for all regions

## Notes

### Competing Interest Statement

The authors have declared no competing interest.

