## Supplementary Figures and Tables for "Multi-region spatial transcriptomics reveals region specific differences in response to amyloid beta (Aβ) plaque induced changes in Alzheimer’s Disease (AD)"

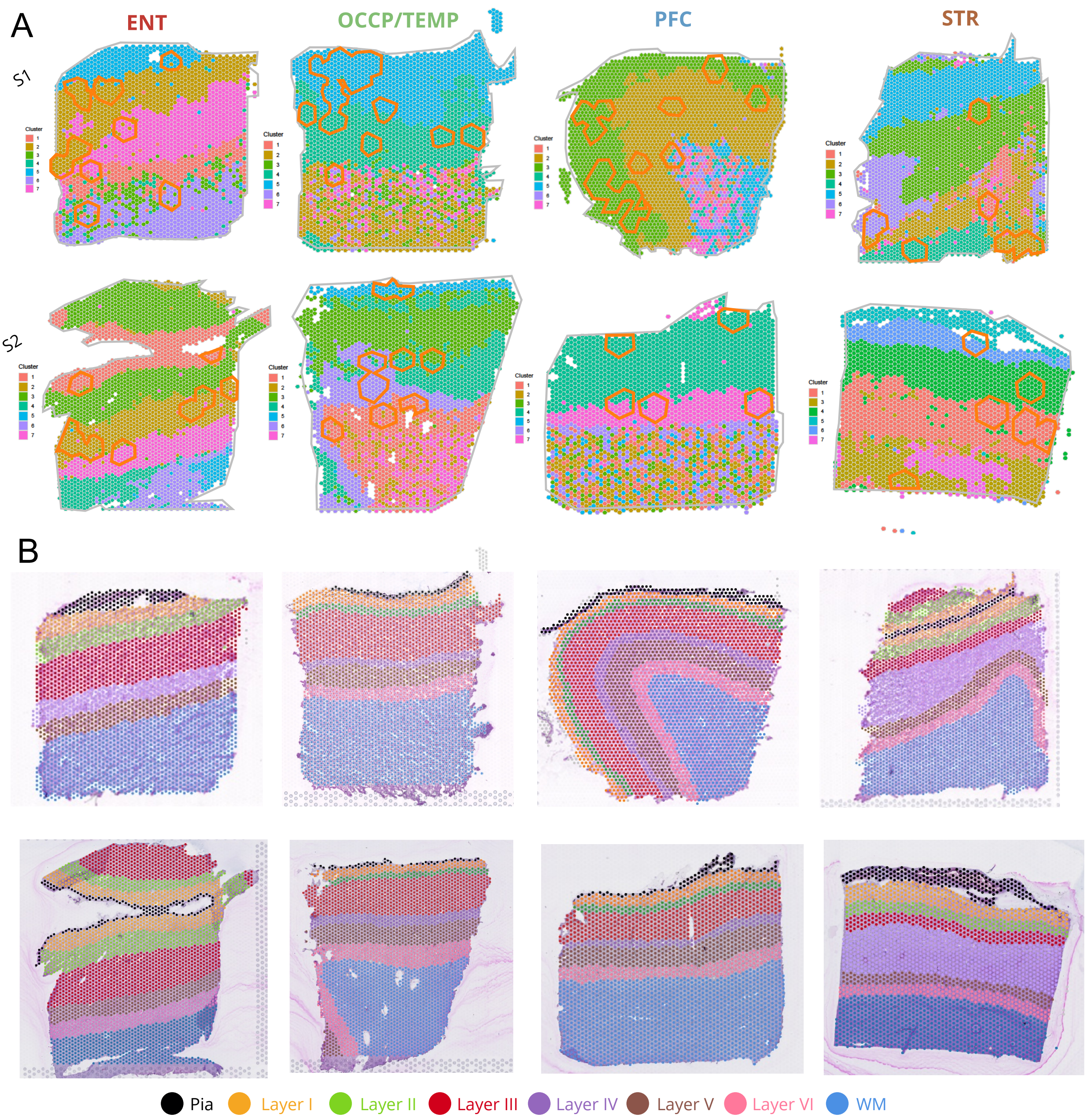


**Supplementary Figure 1**. Clustering and Annotation of Cortex Laminar Layers **A)** BayesSpace Spatial Clustering of each Sample. Each sample was run separately with the number expected clusters set to 7. **B)** Annotation of Laminar Layers. Laminar layers were annotated referencing nuclei density, number of spots determining the cortical thickness and the expected proportions of each layer and the boundaries of BayesSpace clusters.

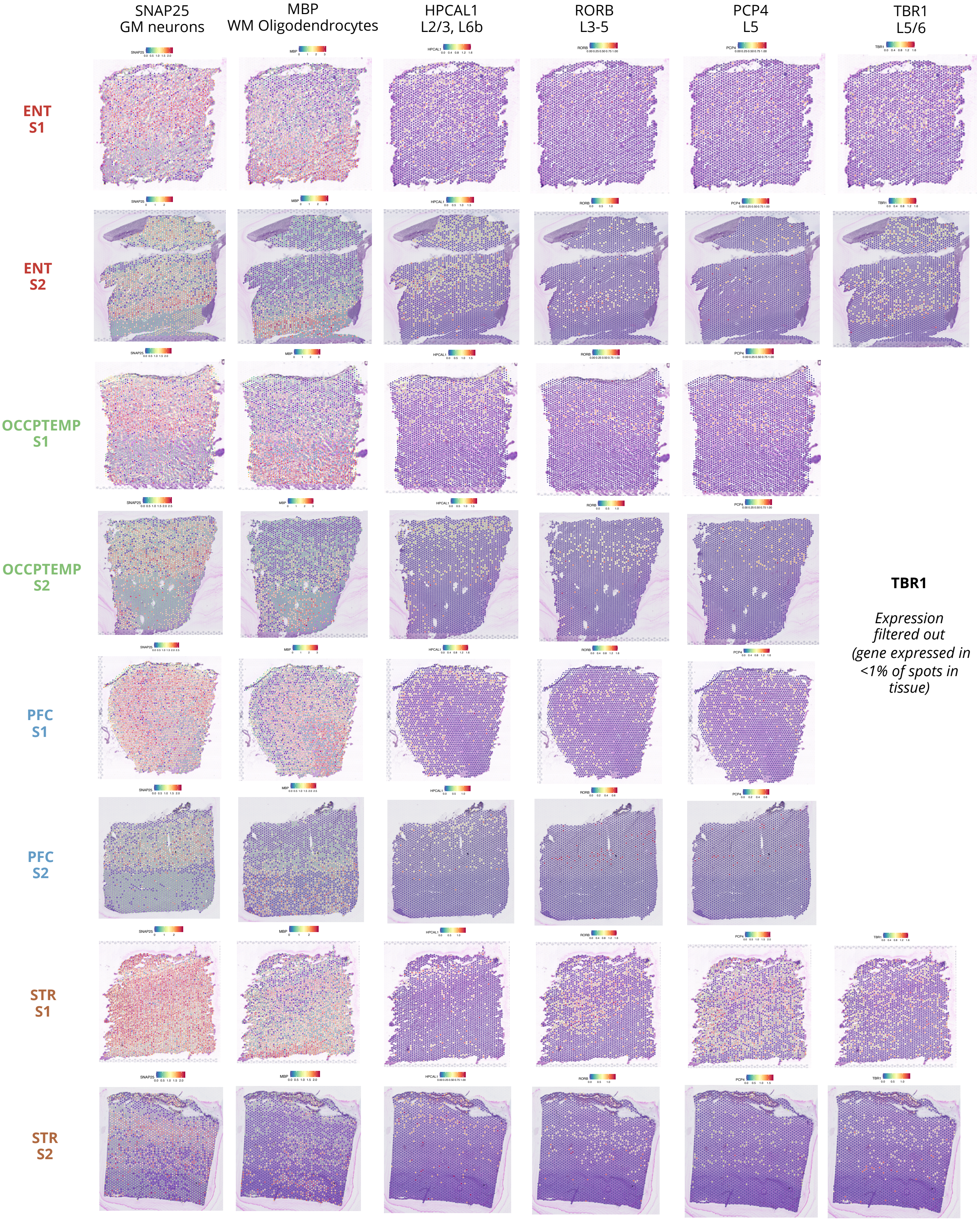


**Supplementary Figure 2.** Spatial Expression of Layer Specific Markers Genes in All 8 Samples. SCTransform-ed gene expression of SNAP25 (GM neurons), MBP (WM Oligodendrocytes/myelin), HPCAL1 (Layer 2/3 and 6b), RORB (Layer 3-5), PCP4 (L5) and TBR1 (L5/6)

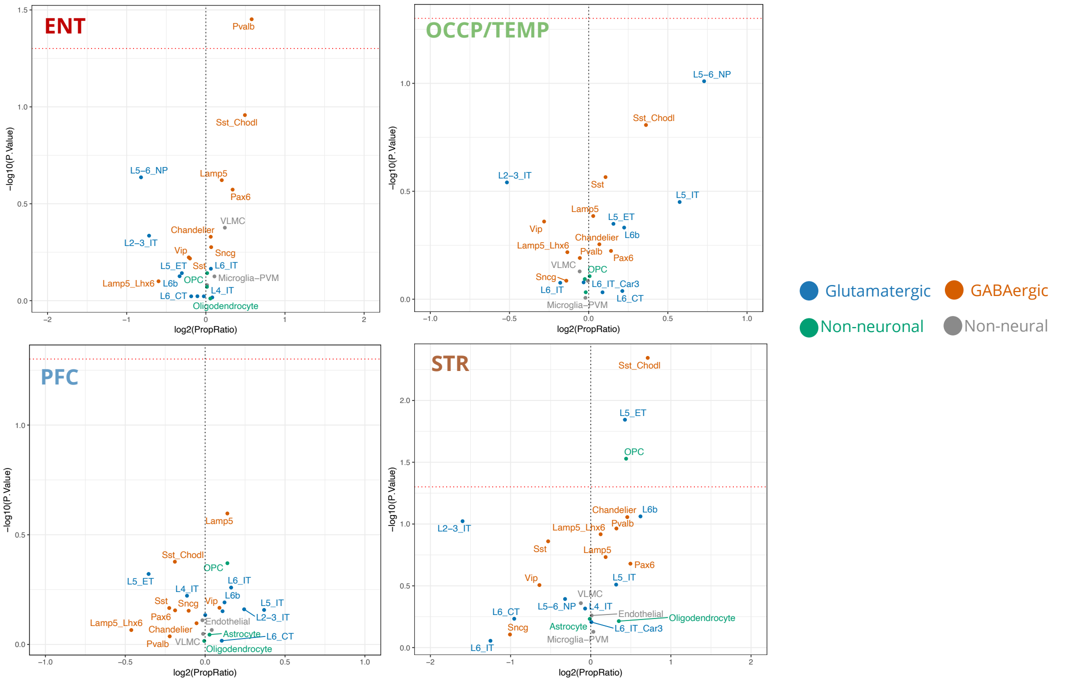


**Supplementary Figure 3**. Differentially Enriched/Depleted Cell Type Proportions the Aβ plaque adjacent spots of all regions vs non-plaque. Log2FC>0 indicating cell types with higher proportions in Aβ plaque adjacent annotated spots, while Log2FC<0 indicate cell types with a lower proportion in Aβ plaque adjacent spots. Spots are colored by cell type category.

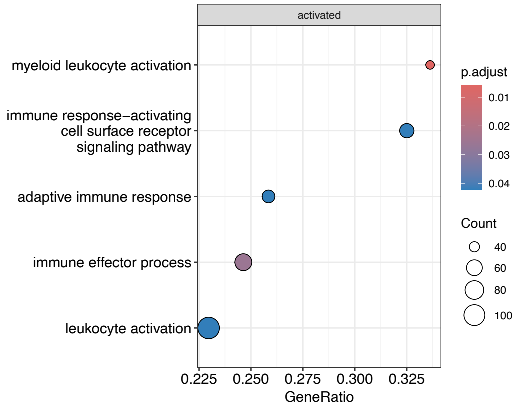


**Supplementary Figure 4**. Dotplot of enriched pathways in DEGs between ENT plaque vs non-plaque ST spots (adjusted p-value <0.05).

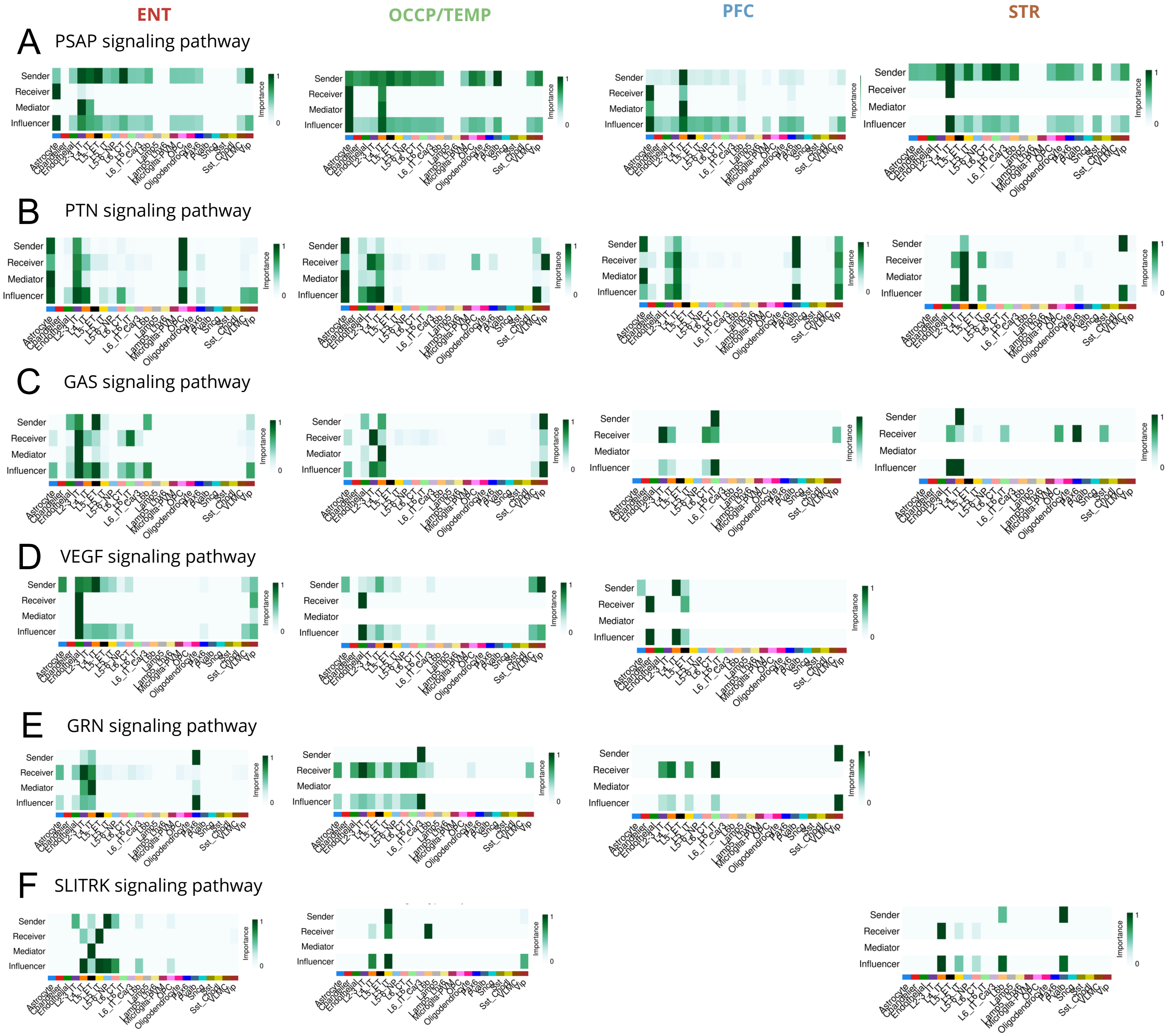


**Supplementary Figure 5**. Cell-cell communication networks for different pathways across the ENT, OCCP/TEMP, PFC and STR regions. Heatmaps of the expected role for each cell type calculated from the network analysis scores. **A)** Prosaposin (PSAP), **B)** Pleiotrophin (PTN) **C)** Growth arrest-specific (GAS) **D)** Vascular endothelial growth factor (VEGF) **E)** Granulin (GRN) and **F)** SLIT- and NTRK-like (SLITRK)

**Suppl Table 1**. Sample Information and Visium Spatial Transcriptomics Raw Quality Control Metrics

| *Sample ID* | *Age* | *Sex* | *APOE Status* | *Thal Phase* | *Braak Stage* | *CERAD score* | *Sections* | *Number of spots under tissue* | *Fraction Reads in Spots Under Tissue* | *Number of reads (million)* | *Total Genes Detected* | *Mean reads per spot* | *Median genes per spot* | *Median UMI counts per spot* |
|---|---|---|---|---|---|---|---|---|---|---|---|---|---|---|
| S1 | 72 | M | APOE3/4 | 4 | III | 2 | ENT | 2,543 | 62.90% | 228 | 18,261 | 89,699 | 755 | 986 |
| | | | | | | | OCCPTEMP | 2,819 | 76.50% | 242 | 18,072 | 85,770 | 709 | 913 |
| | | | | | | | PFC | 3,102 | 83.30% | 229 | 18,062 | 73,711 | 800 | 1,052 |
| | | | | | | | STR | 3,314 | 73.70% | 180 | 18,446 | 54,439 | 977 | 1,381 |
| S2 | 89 | F | APOE3/4 | 4 | III | 1 | ENT | 3,221 | 95.80% | 201 | 18,936 | 62,600 | 1,042 | 1,927 |
| | | | | | | | OCCPTEMP | 2,907 | 95.20% | 210 | 18,201 | 72,082 | 728 | 1,185 |
| | | | | | | | PFC | 2,578 | 90.40% | 186 | 18,081 | 71,967 | 447 | 632 |
| | | | | | | | STR | 2,421 | 85.90% | 217 | 17,942 | 89,611 | 514 | 664 |

**Suppl Table 2**. Mean RCTD weights of 24 cell types for each plaque category (Aβ plaque, Aβ plaque-adjacent, non-plaque)

| | ***ENT_S1__plaque*** | ***ENT_S1__plaque_adjacent*** | ***ENT_S1__non_plaque*** | ***ENT_S2__plaque*** | ***ENT_S2__plaque_adjacent*** | ***ENT_S2__non_plaque*** | ***PFC_S1__plaque*** | ***PFC_S1__plaque_adjacent*** | ***PFC_S1__non_plaque*** | ***PFC_S2__plaque*** | ***PFC_S2__plaque_adjacent*** | ***PFC_S2__non_plaque*** | ***OCCPTEMP_S1__plaque*** | ***OCCPTEMP_S1__plaque_adjacent*** | ***OCCPTEMP_S1__non_plaque*** | ***OCCPTEMP_S2__plaque*** | ***OCCPTEMP_S2__plaque_adjacent*** | ***OCCPTEMP_S2__non_plaque*** | ***STR_S1__plaque*** | ***STR_S1__plaque_adjacent*** | ***STR_S1__non_plaque*** | ***STR_S2__plaque*** | ***STR_S2__plaque_adjacent*** | ***STR_S2__non_plaque*** |
|---|---|---|---|---|---|---|---|---|---|---|---|---|---|---|---|---|---|---|---|---|---|---|---|---|
| ***Astrocyte*** | 0.196081 | 0.190835 | 0.198314 | 0.226229 | 0.215702 | 0.204191 | 0.14651 | 0.148698 | 0.156143 | 0.20376 | 0.184602 | 0.170911 | 0.171814 | 0.16972 | 0.167829 | 0.178942 | 0.160523 | 0.168058 | 0.100071 | 0.076487 | 0.089238 | 0.172054 | 0.180937 | 0.170291 |
| ***Chandelier*** | 0.022078 | 0.015963 | 0.015532 | 0.015154 | 0.014884 | 0.014019 | 0.016053 | 0.018113 | 0.020454 | 0.021476 | 0.016102 | 0.015064 | 0.013382 | 0.016281 | 0.016342 | 0.011455 | 0.01761 | 0.015999 | 0.007874 | 0.031776 | 0.030324 | 0.040163 | 0.045534 | 0.026037 |
| ***Endothelial*** | 0.135015 | 0.135226 | 0.137546 | 0.173724 | 0.156475 | 0.151113 | 0.116663 | 0.116248 | 0.114444 | 0.136256 | 0.138715 | 0.143554 | 0.127297 | 0.125014 | 0.126043 | 0.119826 | 0.130763 | 0.130615 | 0.110422 | 0.120791 | 0.114321 | 0.152121 | 0.147232 | 0.151647 |
| ***L2-3_IT*** | 0.037724 | 0.031717 | 0.026622 | 0.050001 | 0.049612 | 0.107099 | 0.067247 | 0.057329 | 0.041529 | 0.033384 | 0.054699 | 0.053056 | 0.055808 | 0.061921 | 0.058434 | 0.009044 | 0.023819 | 0.064196 | 0.003889 | 0.009811 | 0.028993 | 0.019511 | 0.013002 | 0.040108 |
| ***L4_IT*** | 0.087971 | 0.082338 | 0.087864 | 0.132817 | 0.10711 | 0.091157 | 0.242942 | 0.234894 | 0.201656 | 0.073518 | 0.121724 | 0.184155 | 0.194505 | 0.181831 | 0.157124 | 0.112062 | 0.142381 | 0.174367 | 0.270904 | 0.339213 | 0.367743 | 0.140908 | 0.177145 | 0.174283 |
| ***L5_ET*** | 0.005437 | 0.011777 | 0.013748 | 0.011488 | 0.014093 | 0.01817 | 0.009023 | 0.010449 | 0.015358 | 0.011895 | 0.014636 | 0.016698 | 0.014188 | 0.015976 | 0.015915 | 0.036408 | 0.020847 | 0.017127 | 0.025158 | 0.014551 | 0.008778 | 0.028164 | 0.022088 | 0.018476 |
| ***L5_IT*** | 0.005469 | 0.014723 | 0.021725 | 0.033652 | 0.033023 | 0.026872 | 0.013806 | 0.018207 | 0.027585 | 0.041067 | 0.057316 | 0.03089 | 0.018335 | 0.027529 | 0.034429 | 0.047865 | 0.076946 | 0.035736 | 0.005674 | 0.026753 | 0.020094 | 0.030293 | 0.026068 | 0.022299 |
| ***L5-6_NP*** | 0.003808 | 0.009413 | 0.014811 | 0.003113 | 0.007989 | 0.015889 | 0.011755 | 0.010419 | 0.014074 | 0.028722 | 0.019498 | 0.013666 | 0.009825 | 0.015361 | 0.011596 | 0.020392 | 0.027713 | 0.014395 | 0.029495 | 0.006533 | 0.008227 | 0.015819 | 0.014967 | 0.018613 |
| ***L6_CT*** | 0.0061 | 0.016497 | 0.016352 | 0.004908 | 0.007792 | 0.011256 | 0.004196 | 0.0042 | 0.010088 | 0.00815 | 0.019247 | 0.011728 | 0.006211 | 0.005248 | 0.009842 | 0.028253 | 0.023319 | 0.014812 | 0.005614 | 0.002977 | 0.004878 | 0.002398 | 0.006071 | 0.012654 |
| ***L6_IT*** | 0.011686 | 0.019544 | 0.01529 | 0.004372 | 0.01801 | 0.02064 | 0.005814 | 0.014089 | 0.014397 | 0.019378 | 0.01768 | 0.013983 | 0.012476 | 0.015408 | 0.018811 | 0.027898 | 0.018616 | 0.019725 | 0.007057 | 0.004023 | 0.008108 | 0.011014 | 0.005476 | 0.014498 |
| ***L6_IT_Car3*** | 0.023421 | 0.027699 | 0.023317 | 0.021552 | 0.026503 | 0.035044 | 0.015063 | 0.01621 | 0.029838 | 0.064212 | 0.04938 | 0.035725 | 0.013485 | 0.020092 | 0.027293 | 0.08978 | 0.061052 | 0.049081 | 0.006909 | 0.015481 | 0.017146 | 0.016404 | 0.025703 | 0.023808 |
| ***L6b*** | 0.007324 | 0.015355 | 0.017203 | 0.008269 | 0.006835 | 0.01072 | 0.01341 | 0.00811 | 0.010748 | 0.002762 | 0.014792 | 0.010307 | 0.012163 | 0.010991 | 0.009943 | 0.008827 | 0.013585 | 0.011098 | 0.021535 | 0.018933 | 0.005357 | 0.010869 | 0.005325 | 0.010419 |
| ***Lamp5*** | 0.011668 | 0.017318 | 0.013666 | 0.007217 | 0.007307 | 0.007747 | 0.014846 | 0.014781 | 0.01293 | 0.008121 | 0.009122 | 0.008775 | 0.016399 | 0.012654 | 0.011683 | 0.002632 | 0.007668 | 0.00824 | 0.033857 | 0.023188 | 0.01818 | 0.008449 | 0.010276 | 0.011236 |
| ***Lamp5_Lhx6*** | 0.01157 | 0.005333 | 0.009696 | 0.004082 | 0.005766 | 0.007102 | 0.008701 | 0.009099 | 0.011164 | 0.006289 | 0.003896 | 0.006739 | 0.008602 | 0.009388 | 0.010703 | 0.010476 | 0.006041 | 0.006235 | 0.028376 | 0.012855 | 0.010936 | 0.005099 | 0.009055 | 0.009177 |
| ***Microglia-PVM*** | 0.050517 | 0.049925 | 0.045353 | 0.034483 | 0.038303 | 0.03646 | 0.034612 | 0.032244 | 0.030802 | 0.041664 | 0.037603 | 0.03707 | 0.033323 | 0.035372 | 0.039212 | 0.038943 | 0.036527 | 0.033718 | 0.016152 | 0.018418 | 0.025741 | 0.049026 | 0.045255 | 0.036493 |
| ***Oligodendrocyte*** | 0.143163 | 0.129647 | 0.124893 | 0.053617 | 0.051543 | 0.049319 | 0.100936 | 0.100485 | 0.112755 | 0.08714 | 0.082837 | 0.071269 | 0.095525 | 0.096789 | 0.099572 | 0.054392 | 0.057961 | 0.057179 | 0.074999 | 0.089695 | 0.074155 | 0.105756 | 0.099685 | 0.074368 |
| ***OPC*** | 0.021841 | 0.022305 | 0.020764 | 0.015936 | 0.022351 | 0.02338 | 0.01746 | 0.017997 | 0.019262 | 0.010066 | 0.022008 | 0.01706 | 0.023005 | 0.020743 | 0.02138 | 0.013687 | 0.021573 | 0.020762 | 0.030632 | 0.025073 | 0.01441 | 0.024952 | 0.021515 | 0.0199 |
| ***Pax6*** | 0.057771 | 0.054315 | 0.050135 | 0.046401 | 0.052162 | 0.03407 | 0.027668 | 0.025508 | 0.02252 | 0.025715 | 0.01698 | 0.025884 | 0.036482 | 0.028312 | 0.02606 | 0.026405 | 0.031904 | 0.028549 | 0.044394 | 0.048779 | 0.027494 | 0.020901 | 0.019738 | 0.021114 |
| ***Pvalb*** | 0.02203 | 0.01614 | 0.010397 | 0.00872 | 0.013489 | 0.009435 | 0.010764 | 0.011989 | 0.011172 | 0.007651 | 0.0052 | 0.008874 | 0.008696 | 0.010372 | 0.010823 | 0.007826 | 0.009935 | 0.010285 | 0.030537 | 0.023433 | 0.020881 | 0.011367 | 0.01985 | 0.01378 |
| ***Sncg*** | 0.011206 | 0.013906 | 0.013601 | 0.009634 | 0.012772 | 0.011857 | 0.006846 | 0.009927 | 0.012052 | 0.025911 | 0.009382 | 0.008686 | 0.014462 | 0.010415 | 0.012716 | 0.021517 | 0.009498 | 0.009242 | 5.52E-05 | 0.002724 | 0.011407 | 0.009582 | 0.007527 | 0.009198 |
| ***Sst*** | 0.006958 | 0.006513 | 0.00967 | 0.003619 | 0.008082 | 0.0071 | 0.003125 | 0.007467 | 0.009763 | 0.003961 | 0.006405 | 0.006447 | 0.010349 | 0.010156 | 0.010791 | 0.008381 | 0.008041 | 0.006113 | 0.02866 | 0.000649 | 0.007718 | 0.015907 | 0.010307 | 0.008115 |
| ***Sst_Chodl*** | 0.010474 | 0.012849 | 0.009156 | 0.004178 | 0.005476 | 0.003856 | 0.004553 | 0.004555 | 0.005305 | 0.00772 | 0.003534 | 0.00392 | 0.005588 | 0.005996 | 0.00516 | 0.008811 | 0.00627 | 0.004386 | 5.52E-05 | 0.00656 | 0.003108 | 0.005767 | 0.006665 | 0.00497 |
| ***VLMC*** | 0.09082 | 0.072974 | 0.07539 | 0.096295 | 0.098158 | 0.069467 | 0.0689 | 0.071377 | 0.062008 | 0.10803 | 0.075217 | 0.085895 | 0.064816 | 0.059795 | 0.058295 | 0.095848 | 0.069379 | 0.076079 | 0.105221 | 0.064926 | 0.056497 | 0.079631 | 0.064984 | 0.084975 |
| ***Vip*** | 0.019869 | 0.027689 | 0.028954 | 0.030538 | 0.026564 | 0.034034 | 0.039109 | 0.037605 | 0.033954 | 0.02315 | 0.019426 | 0.019644 | 0.033265 | 0.034636 | 0.040005 | 0.020328 | 0.018027 | 0.024001 | 0.01246 | 0.016371 | 0.026266 | 0.023847 | 0.015594 | 0.023543 |
